# RootDigger: a root placement program for phylogenetic trees

**DOI:** 10.1101/2020.02.13.935304

**Authors:** Ben Bettisworth, Alexandros Stamatakis

## Abstract

**Summary:** In phylogenetic analysis, it is common to infer unrooted trees. Thus, it is unknown which node is the most recent common ancestor of all the taxa in the phylogeny. However, knowing the root location is desirable for downstream analyses and interpretation. There exist several methods to recover a root, such as midpoint rooting or rooting the tree at an outgroup. Non-reversible Markov models can also be used to compute the likelihood of a potential root position. We present a software called `RootDigger` which uses a non-reversible Markov model to compute the most likely root location on a given tree and to infer a confidence value for each possible root placement.

**Availability and implementation:** `RootDigger` is available under the MIT licence at https://github.com/computations/root_digger

## 1 Introduction

Most phylogenetic inference tools [7, 10, 9] return unrooted trees because they typically implement time-reversible nucleotide substitution models [2] as they yield the phylogenetic inference problem computationally tractable. However, time-reversible models are incapable of identifying root. But, having a rooted phylogeny is often required for downstream analyses and interpretation of results.

A given phylogeny can be rooted via an outgroup or a molecular clock analysis. Unfortunately, both approaches exhibit their own problems and pitfalls [4, 6, 1]. Alternatively one can use a non-reversible model as, under such a model, the root placement affects the likelihood [14]. It can thus be used to determine the most likely root placement. However, using a non-reversible model during tree search increases the computational complexity *per tree* by a factor of 𝒪(*n*), where *n* is the number of taxa. Alternatively, one can infer a root on a given unrooted tree via a non-reversible model as this requires substantially less computational effort.

## 2 The RootDigger Software

`RootDigger` infers a rooting on an existing phylogeny using a non-reversible model. The input is a multiple sequence alignment (MSA) and a give phylogeny with branch lenghts in expected substitutions per site. `RootDigger` then finds the most likely root location on the given phylogeny by calculating the likelihood of a rooting under the non-reversible UNREST model [13] with a user specified number of discrete Γ rate categories and, optionally, a proportion of invariant sites estimate. `RootDigger` implements fast and a slow root finding modes, called *Search* mode and *Exhaustive* mode, respectively. The search mode finds the most likely root quickly via appropriate heuristics. To do this, `RootDigger` conducts a user-specified number of independent root searches with random initial roots in order to avoid local optima. In contrast, the Exhaustive mode computes the likelihood and the uncertainty (via the likelihood weight ratio (LWR) [11]) of rooting the tree on each branch.

Each search mode can be executed under an optional early stop mode which terminates the search if two successive root placements are on the same branch *and* their root positions along this branch are “sufficiently” close. Early stopping improves run times by up to a factor of 1.7 on some empirical datasets while not substantially decreasing root placement accuracy. This does, however, render likelihood-based comparisons with other tools invalid as it only computes an approximate likelihood.

## 3 Experiments and Results

To validate `RootDigger` and investigate the effects of early stopping, we conducted experiments on simulated *and* empirical data.

### Simulations

Simulated data tests were conducted to validate the software and compare it against IQ-TREE version 1.6.11 [7]. We simulated trees and alignments using ETE3 [5] and INDELible[3]. IQ-TREE and `RootDigger` were given the same model options for all runs (see supplement for details).

To simulate the operation of `RootDigger` we constrained IQ-TREE by a fully resolved unrooted tree as, in this case, it will simply root the tree. A notable difference between the results of IQ-TREE and `RootDigger` is that IQ-TREE will also optimize all branch lengths for every candidate tree. Unfortunately, there was no option to disable this at the time of writing the paper (from personal communication). To compensate, we have attempted to adjust the IQ-TREE running time by removing the time spent in branch length optimization (see supplement for details). The additional computational effort for branch length optimization in IQ-Tree renders our execution time comparisons useful for reference only, and should be interpreted with care.

In our experiments, we varied the number of MSA sites (1000 and 8000) and the number of taxa (10 and 50). The results as well as the execution times, are shown in figure 1. Additional simulations are presented in the supplement.

**Figure 1:**
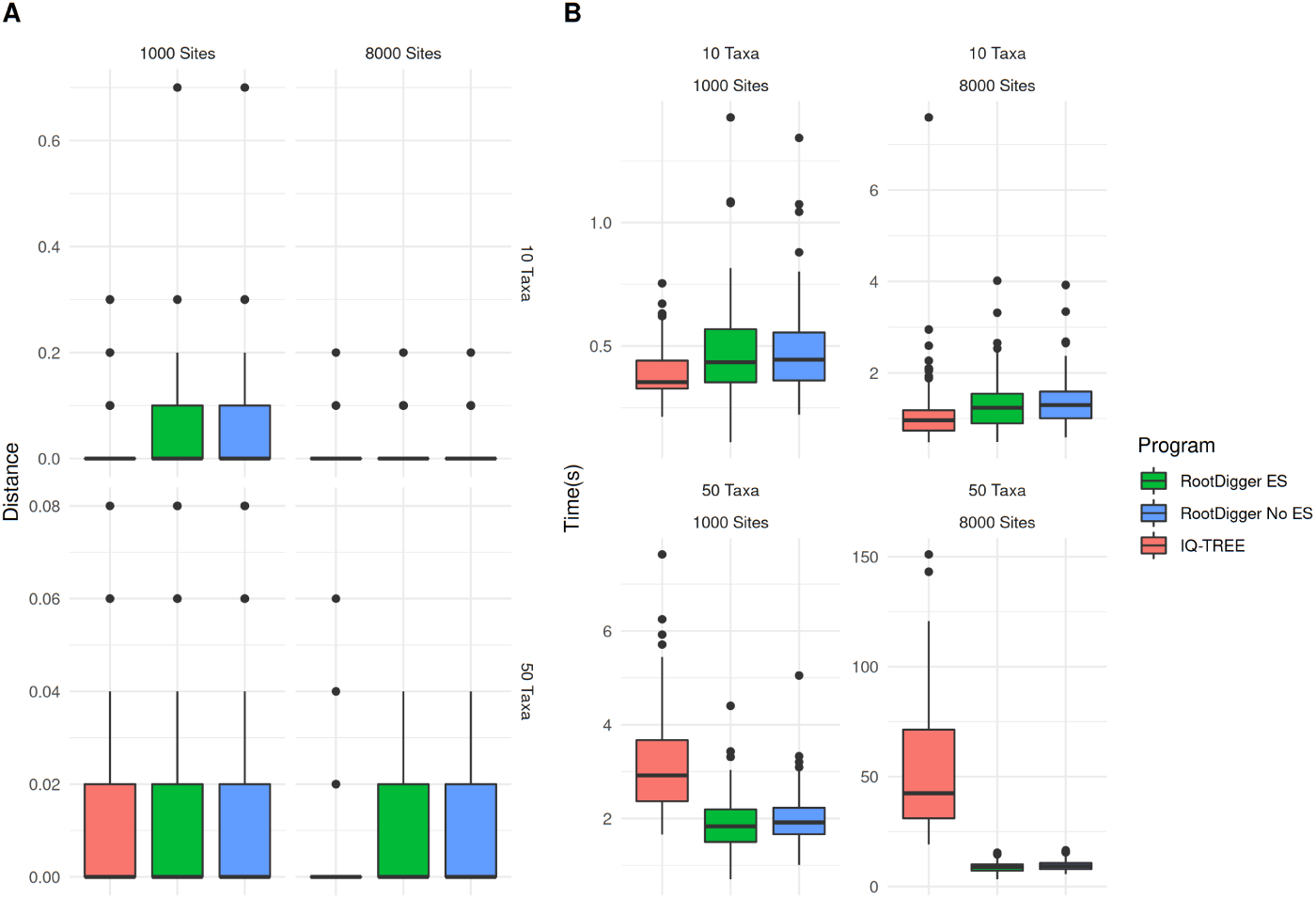
Comparison of `RootDigger` in search mode and IQ-TREE on result error (A) and execution time (B). The distance in (A) is the topological distance from the inferred root to the true root normalized by the number of taxa on the tree. ES stands for Early Stop, and indicates that early stopping was used. The execution time for IQ-TREE has been discounted.

### Empirical Data

The empirical datasets were chosen from TreeBASE [8, 12] to include an existing, strongly supported outgroup and are listed in Table 1. To ensure that the branch lengths are specified in substitutions per site, we initially re-optimized branch lengths using RAxML-NG version `0.9.0git` under the model used in the original publication or GTR+G4 if it was not known. Then, we ran `RootDigger` on each datasets in exhaustive mode to obtain likelihood weight ratios (LWR) for placing the root into each branch (see supplement for LWR annotated trees).

**Table 1:** Table of empirical datasets used for validation and results.

| Dataset | Taxa | Sites | RD D<sup>a b</sup> | IQ D<sup>a</sup> | Search<sup>a b</sup> | Exhaus | IQ<sup>a d</sup> |
| --- | --- | --- | --- | --- | --- | --- | --- |
| DS1<sup>c</sup> | 35 | 864kb | 0.000 |  | 13m | 1.1h |  |
| DS2<sup>c</sup> | 35 | 1296kb | 0.143 |  | 36m | 3.3h |  |
| DS3<sup>c</sup> | 245 | 5kb | 0.000 | 0.004 | 3.1m | .86h | 9.5m |
| DS4<sup>c</sup> | 200 | 6kb | 0.075 | 0.055 | 0.5m | .27h | 0.8m |
| DS5<sup>c</sup> | 33 | 1097kb | 0.000 |  | 5.5m | 1.1h |  |
| DS6<sup>c</sup> | 34 | 12kb | 0.034 | 0.088 | 0.3m | .04h | 1.8m |

<sup>a</sup> Averaged over 100 independent executions
<sup>b</sup> In early stop mode
<sup>c</sup> See supplement for more information
<sup>d</sup> With time spent optimizing branch lengths removed as measured with the profiling tool `perf`.

The results for empirical datasets are given in Table 1. The evaluation uses topological distance from the root inferred by `RootDigger` or IQ-TREE to the “true”, or the root indicated by the outgroup, root, normalized by the number of taxa in the tree. We also report execution times for Search mode, Exhaustive mode, and IQ-TREE. Search mode was conducted *with* early stop mode enable, and Exhaustive mode was conducted *without*. Times reported for IQ-TREE are adjusted down to compensate for the extra work required for branch length optimization. In our experiments, the maximum proportion of time spent in branch length optimizations was 56.9

### Effect of early stopping

We investigated the effect of the early stopping criterion on the final LWR results in exhaustive mode by executing `RootDigger` on all empirical datasets with early stopping enabled and disabled. For most runs, the results with and without early stopping showed no meaningful (difference in LWR less than 0.000001) difference. The dataset that showed the largest difference in LWR is DS6 (see supplement for details). `RootDigger` is about 1.7 times faster with early stopping enabled.

## 4 Discussion

Compared to IQ-TREE, `RootDigger` performs competitively, as can be seen in both sides of Figure 1. The results on simulations are mixed, with IQ-TREE performing slightly better in terms of root placement in most scenarios. We see 2 possible reasons: First, the branch length optimization that IQ-TREE performs induces a slight accuracy advantage; or the longer search time allows for IQ-TREE to be more thorough.

`RootDigger` was faster on all but the smallest datasets. On the smallest datasets, IQ-TREE and `RootDigger` performed about equal in terms of execution time. Again, it should be noted that IQ-TREE *does* optimize branch lengths for *every* candidate tree, while `RootDigger` does not. Thus, `RootDigger` being faster than IQ-TREE is expected. Nonetheless, IQ-TREE is the only tool that will find the same results as `RootDigger` with comparable methodology. Hence, the execution times are only presented here for context.

For empirical datasets `RootDigger` performed well, correctly identifying the root for 3 of the 6 datasets, which is consistent with the results from simulations. The time required to analyze these datasets in exhaustive mode ranged from approximately 23 minutes (DS6) to 3.3 hours (DS2) with early stopping disabled.

Given the computational cost of obtaining a runtime sample for analyzing the largest datasets with IQ-TREE, we chose not to compare with `RootDigger` on every empirical dataset. Instead, we report a performance comparison on a subset of datasets: DS3, DS4, and DS6.

Qualitatively, `RootDigger` offers other advantages: multiple modes of inference (search mode and exhaustive mode), as well as the ability to compute LWRs for all putative root positions on a given tree.

Furthermore, the accuracy of root placement of `RootDigger` is nearly the same as IQ-TREE when using and not using the early stop heuristic, as seen in Figure 1. This indicates that our heuristics, explicitly, not optimizing the branch lengths and stopping early, were effective in reducing execution times while not impacting accuracy. The largest observed difference on empirical datasets between using early stop mode and full optimization is negligent (see supplementary material). Finally, `RootDigger` outperforms IQ-TREE in head to head benchmarks on empirical data (see Table 1).

The computational improvement of the early stop mode on simulated datasets is less prevalent than on empirical datasets. Early stop mode on simulated datasets performed at negligibly better (the largest improvement is approximately 2%) compared without early stop mode. This is almost certainly due to the noise free nature of simulated data. The lack of additional noise causes the area around likelihood peaks to be very steep, which makes them particularly amenable to optimize using traditional routines.

In conclusion, `RootDigger` is an easy to use and competitive software package to root an existing unrooted tree. In addition, we have plans to develop a web service for `RootDigger`.

## Supporting information

Supplemental Paper

## Acknowledgments and Funding

The authors gratefully acknowledge the support of the Klaus Tschira Foundation. This project has received funding from the European Union’s Horizon 2020 research and innovation programme under the Marie Sklodowska-Curie grant agreement No 764840.

## Notes

#### Summary of Updates

Tables and figures were updated with an adjusted execution times for IQ-TREE in an attempt to be more fair to the competition.

