## Supplemental Paper for "RootDigger: a root placement program for phylogenetic trees"

March 3, 2020

#### 1 Online Supplement Structure

In Section 2 we present additional information relevant to the process of rooting an existing tree, as well as the background to `RootDigger`. Section 3 gives a detailed description of the exact process that `RootDigger` uses to infer a root. Section 4 describes dependencies availability and usage for `RootDigger`. Section 5 goes into more detail about the experiments and results of both simulations and empirical tests. Finally Section 6 contains discussion about the results in the context of other tools and previous work.

#### 2 Background

As mentioned in the paper, when attempting to root an existing phylogenetic tree, there are two common methods: including an outgroup, or molecular clock analysis. We briefly discussed these, along with a longer discussion of non-reversible models. However, we feel that a stronger justification is required. Therefore, we present a more general overview of methods used to root a tree, and discuss the problems associated with each of them in more detail. These are categorized into: methods which use additional topological information not present in the sequence data; methods which utilize some variation of the molecular clock hypothesis; and methods which utilize a non-reversible model.

Methods that use additional topological information take advantage of prior knowledge about the world, which is not present in the, generally molecular, data that is used to infer the tree. In particular, knowledge about species which are distantly related can be used to include a so-called outgroup to phylogenetic analysis. This outgroup can then be used to place the root on the tree, as the most recent common ancestor of the ingroup and the outgroup must be the root of the tree.

There are challenges to including an outgroup to an analysis. Gatsey et. al. [8] showed how adding a single taxon to an analysis can substantially change the resulting tree topology, even for the taxa which were already present in the analysis (i.e., the ingroup). Holland [10] investigates this phenomenon in simulations, and finds that outgroups that affect or alter the topology of ingroups are common.

Alternatively, molecular clock analysis can be used to place a root without prior topological knowledge [28]. The molecular clock hypothesis supposes that base substitution "ticks" at a stochastically constant rate. Using this supposition, or some variant thereof, a likely location can be inferred for the root on an existing phylogenetic tree. A simple version of this is midpoint rooting, which relies on a constant molecular clock assumption.

Molecular clock analyses exhibit their own difficulties. In particular, the clock does not generally "tick" at a constant rate over the tree [15, 23]. Relaxed clock models exist which can alleviate this problem, but are not always successful at correctly identifying the root as shown in [1] and come with their own set of inference errors.

The final method that can place a root on a tree is to perform the phylogenetic analysis using a non-reversible model. When using a non-reversible model, the direction of time affects the likelihood of the tree [29]. Using this, the most likely location on the tree for the root can be found. Several software packages are able to infer a phylogenetic tree under a non-reversible model, and as a by product also identify a root [18, 22].

Non-reversible models for phylogenetic trees come in many forms. For example, accounting for duplication, transfer, and loss events yields a non-reversible model [17]. In particular, duplication events have been used for rooting trees [4]. Another method, the one primarily used in this work, is to eliminate the reversibility assumption of standard character (e.g., nucleotide or amino acid) substitution models. Unfortunately, eliminating this assumption significantly increases the computational effort required to find a good (high likelihood) phylogenetic tree. This is due to the loss of the pulley principle [5], which allows phylogenetic inference tools to ignore root placement during tree inference. Therefore, by adopting a non-reversible model, the location of a root on a phylogenetic tree affects the likelihood of that tree.

As the location of the root affects the likelihood of the tree, when using standard tree search techniques all possible rootings would need to be evaluated for each tree in order to find the rooting with the highest likelihood. In the worst case, this increases the work *per tree* by a factor of $\mathcal{O}(n)$ where $n$ is the number of taxa present in the dataset. Therefore, eliminating the reversibility assumption drastically increases the computational effort required to infer a tree. Hence, standard inference tools choose to adopt the reversibility assumption, as phylogentic tree inference would be computationally intractable otherwise.

As an alternative to the computationally expensive process of inferring a tree with a root, an unrooted tree which has already been inferred under a reversible model can be evaluated a posteriori for possible root locations under a non-reversible model. This requires less computational effort, as it skips the expensive step of looking for “good” rootings in intermediate trees during the tree search. With this method, we can find the most likely root location for a given phylogenetic tree. Even this method still has numerical challenges, as previous research suggests the likelihood function for rooting a phylogenetic tree may exhibit several local maxima [11], although we did not find this to be a major issue in our experiments (see the discussion).

We implemented the open source software tool `RootDigger` which uses a non-reversible model of character substitution to infer a root on an already inferred tree. The inputs to our tool are a multiple sequence alignment (MSA) and a phylogenetic tree. `RootDigger` then yields a rooted tree. `RootDigger` implements fast and a slow root finding modes, called Search mode and Exhaustive mode, respectively. The search mode simply finds the most likely root quickly via appropriate heuristics. The exhaustive mode evaluates the likelihood of placing the root into every branch of the given tree, and reports the likelihood weight ratio [24] for placing a root on that branch for every branch on the tree. In addition to the two search modes, there is an optional early stop mode, which can be combined with either root finding mode. In this early stop mode, the search will terminate if the root placement is nearly the same twice in a row. This is to say, if the location of the best root position is on the same branch as in the previous iteration *and* the value inferred for the root position is sufficiently close the current iteration terminates. While the early stop optimization does improve rooting times substantially (approximately a factor of 1.7 on some empirical datasets), the likelihood of each root placement will not be fully optimized. In practice, this does not substantially affect the final root placement, but it does render comparison of the likelihood with results from other tools invalid.

### 3 Methods

In the following we provide a more detailed technical description of the `RootDigger` algorithm and implementation.

The input to `RootDigger` is a MSA and a phylogenetic tree with branch lengths in expected mean substitutions per site. `RootDigger` then uses the tree and branch lengths to find the most likely root location by calculating the likelihood of a root location under a non-reversible model of DNA substitution (specifically, UNREST [27] with a user specified number of $\Gamma$ discrete rate categories). The optimal position of the root along a specific branc hof length $t$ is calculated by splitting the given branch in two with resulting branch lengths $\beta t$ and $(1 - \beta)t$, with $0 \leq \beta \leq 1.0$. We then find the maximum likelihood value of $\beta$, and report the likelihood for the given branch as the likelihood of the root location. By formulating the problem this way, we can use single parameter optimization techniques such as Brent's Method, which are computationally more efficient compared to multi-parameter optimization routines such as BFGS. Note that we specifically selected Brent's Method instead of using Newton's Method, because it does not require the calculation of the second derivative to optimize the function. We computed an analytical version of the second derivative, but the library which does the likelihood computations that we use does not support calculating some of the non-logged likelihoods. The effort to rewrite the functions that would compute these values deemed to be too much relative to the savings. Nonetheless, in principle, the computation of the second derivative of the likelihood is feasible and could be implemented.

A potential problem of Brent's and analogous methods is that they find extrema by identifying roots for the derivative of the objective function. In order to find maxima, though, it is required that the objective function's value is evaluated, as a root of the derivative could correspond to a minima. In addition, Brent's method will fail to find all extrema. To alleviate this, we need to search for bracketing windows that can be used to safely find extrema. Unfortunately, we are not aware of a general method for finding such bracketing windows, so a recursive method is employed, were the search range is bisected and adequately searched for appropriate windows. Appropriate here means that the sign of the function in question has opposite signs at the endpoints of the window.

As mentioned in the main text, `RootDigger` has 2 modes of operation. These modes will be discussed individually, starting with search mode:

1. Initialize numerical model parameters:
    - $\alpha$-shape parameter for discrete $\Gamma$ rates to 1.0 (if applicable),
    - Character substitution rates to uniform random values according to $U(0.0001, 1.0)$,
    - Base frequencies to uniform random values according to $U(0.0001, 1.0)$, which are then normalized such that the sum is equal to 1.
2. Choose starting roots by best log-likelihood at each branch midpoint (default 5%).
3. For each starting root:
    (a) Optimize model parameters
        - $\alpha$-shape parameter for $\Gamma$ distributed rates (if applicable),
        - Character substitution rates,
        - Base Frequencies.

    (b) Find the best root location for the current model
        i. Create a list of high likelihood root locations evaluated at the midpoint of every branch.
        ii. For the top roots (default 5%), optimize the root location along their specific branch.

    (c) Repeat from $3(a)$ until a stopping condition is met:
        - The difference between likelihoods between this iteration and the previous iteration is sufficiently small (below user defined parameter `atol`),

- If early stopping is enabled, the new root location is sufficiently close to the old root location by distance along the branch (below user defined parameter `brtol`) or,
- More than 500 iterations have passed.

4. Report the best found root, along with its log-likelihood

During the search, we re-estimate both the shape parameter $\alpha$ and the base frequencies every iteration to ensure a good likelihood, and because the cost of optimizing these parameters is small (approximately 0.4% and 10% of run time for $\alpha$ and base frequencies respectively). Furthermore, because we use a non-reversible general substitution matrix, the base frequencies might not be stable across every branch of the tree. Therefore, to ensure a good fit, we need to optimize the base frequencies every time. The process for exhaustive mode is analogous, with the core optimization routines being shared with search mode. The major difference is that all branches are considered:

1. For every branch on the tree:
   (a) Place root at current branch.
   (b) Initialize numerical parameters:
      - $\alpha$-shape parameter for $\Gamma$ distributed rates (if applicable),
      - Character substitution rates,
      - Base Frequencies.
   (c) Optimize model parameters
      - $\alpha$-shape parameter for $\Gamma$ (if applicable),
      - Character substitution rates,
      - Base Frequencies.
   (d) Repeat from $1(c)$ until a stopping condition is met:
      - The difference between likelihoods between this iteration and the previous iteration is sufficiently small (below `atol`) or,

    - If early stopping is enabled, the new root location is sufficiently close to the old root location by distance along the branch (below `brtol`).
    - More than 500 iterations have passed.

2. Report the tree with annotations for every branch:
    - The root position along the branch,
    - The log-likelihood,
    - and the Likelihood Weight Ratio [24].

We re-initialize the initial model parameters in every iteration (from (3) in search mode and (1) in exhaustive mode) to avoid the numerous local minima, as discussed in [11]. In both modes, there is an upper limit to the number of iterations of 500. In empirical and simulated datasets this limit has never been met, and only exists to ensure that the program will eventually halt.

### 4 The Software

In order to implement both likelihood computations and non-reversible models, `RootDigger` has three major dependencies: GNU Scientific Library (GSL) [9], the Phylogenetic Likelihood Library (LibPLL) [7], and L-BFGS-B [30]. GSL is used for the decomposition of a non-symmetric rate matrix, LibPLL is used for efficient likelihood calculations, and L-BFGS-B is used for multi-parameter optimizations.

Usage of `RootDigger` is straight-forward. All that is required is a tree in newick format, and a MSA in either PHYLIP or FASTA format. `RootDigger` is open source, released under the MIT license, and written in C++. The code, documentation, test suite, benchmarks, experiment scripts, datasets, this paper, as well as any modifications to existing libraries can be found at `github.com/computations/root_digger`.

### 5 Experiments and Results

To validate `RootDigger`, we conducted several experiments on both simulated and empirical data. Furthermore, we also used Likelihood Weight Ratios (LWR) [24] to asses the confidence of root placements on empirical datasets. Finally, we investigated the effects of the early stop mode on the final results.

#### 5.1 Simulations

Testing with simulated data was done to both validate the software and to compare against IQ-TREE [16]. We created a pipeline to

1. Generate a random tree with ETE3 [12] and random model parameters.
2. Simulate an MSA with indelible [6]
3. Execute `RootDigger` and IQ-TREE [16] with the simulated MSA, given the generated random tree.
4. Repeat from (2) for a total of 100 iterations
5. Compute comparisons
    (a) Calculated rooted RF distance with ETE3 [21]
    (b) Mapped root placement onto original tree with the true root.

Both IQ-TREE and `RootDigger` were given the same model options for all the runs. `RootDigger` was executed with the arguments

```
rd --msa <MSA FILE> --tree <TREE FILE>
```

By default `RootDigger` uses no rate categories, and currently only supports the UNREST model [27]. IQ-TREE was executed with the arguments

```
iqtree -m 12.12 -s <MSA FILE> -g <TREE FILE>
```

The `-m 12.12` argument to IQ-TREE specifies that the UNREST model should be used [26] and the `-g <TREE FILE>` option constrains the tree search to the given user tree. When given an fully resolved unrooted tree, this has the effect of rooting the tree. We used this option to simulate the operation of `RootDigger`. A notable difference between results of IQ-TREE and `RootDigger` is that IQ-TREE also will optimize all branch lengths, and there is no option at the time of writing to turn off this feature. For that reason, comparison of results between the two tools will be done using topological distance. Furthermore, the added computational effort involved in branch length optimization renders execution time comparisons useful for context only. This is to say, we include execution time comparisons for reference only.

For all runs, the UNREST model was used. Furthermore, we vary two additional parameters used for the size of the data in the simulations: number of MSA sites and taxa. In total, we ran 9 simulated trials with MSA sizes of 1000, 4000, and 8000 sites as well as tree sizes of 10, 50, and 100 taxa. The results from these experiment, as well as the execution times, are shown in figure 1.

In order to compensate for IQ-TREE requiring more work, we measured the time spent for branch length optimization, and discounted the time reported. We did this by running building IQ-TREE with the build type `Profile`, and then running IQ-TREE on the simulated datasets with `perf`. We recorded call graph information with `perf`, and used that to estimate the proportion of time spent optimizing branch lengths. This was done by recording the proportion of samples in the `computeNonrevLikelihoodDervSIMD` function, as this function contains the almost all of the workload associated with optimizing branch lengths. There is additional work associated with branch length optimization, but it accounts for a small portion of total run time. This additional work in IQ-TREE is each NR step (typically $\approx 5$ flops), and the work to iterate over all branches. In total, we estimate this extra work to be at most $\approx$ 2.5% of run time, and so the results are not significantly different by not including it in the discounting. Using the proportion of samples $p$, we discounted by the run time of IQ-TREE by $1 - p$ which represents the time required to find the tree, without branch length optimization. Note that this will not discount extra work induced by changing branch lengths, such as having to re-optimize other model parameters. These adjusted times are only in the main paper. In this supplement we present the unadjusted times in plots and figures

#### 5.2 Empirical Data

In addition to simulated data, we conducted tests with empirical data. The datasets used are described in Table 1. The empirical datasets were chosen from TreeBASE [19, 25] to include an

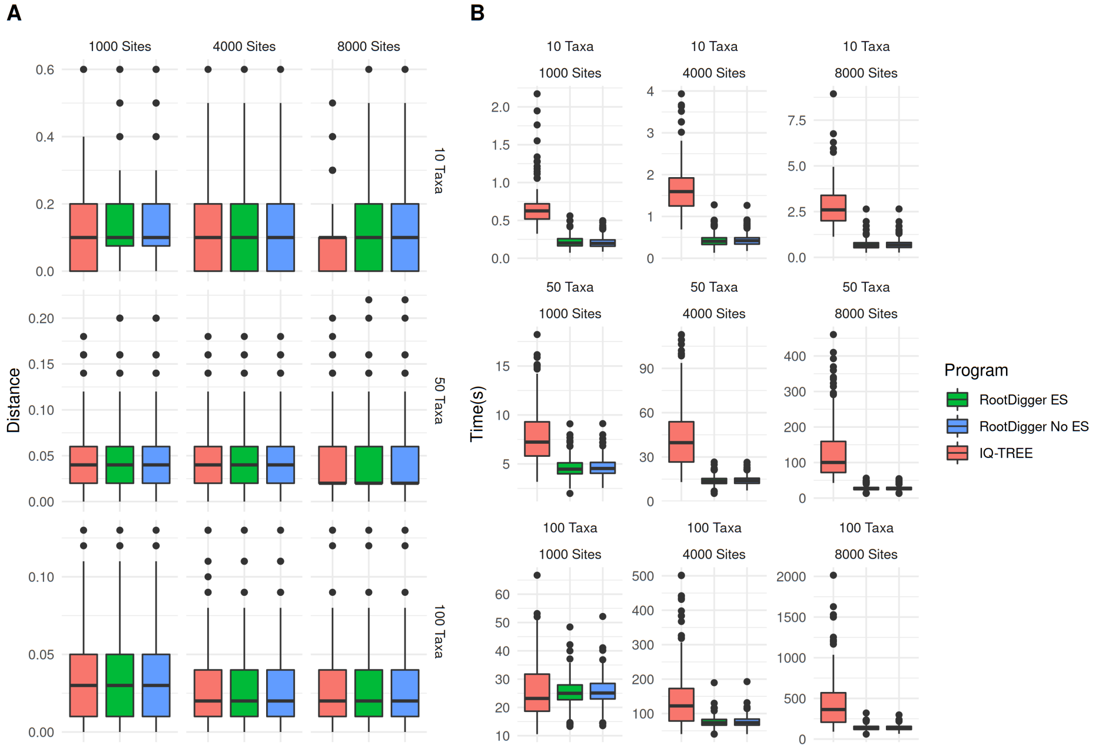


Figure 1: Comparison of `RootDigger` and IQ-TREE on result error (A) and execution time (not discounted) (B). The distance in (A) is the topological distance from the inferred root to the true root normalized by the number of taxa on the tree.

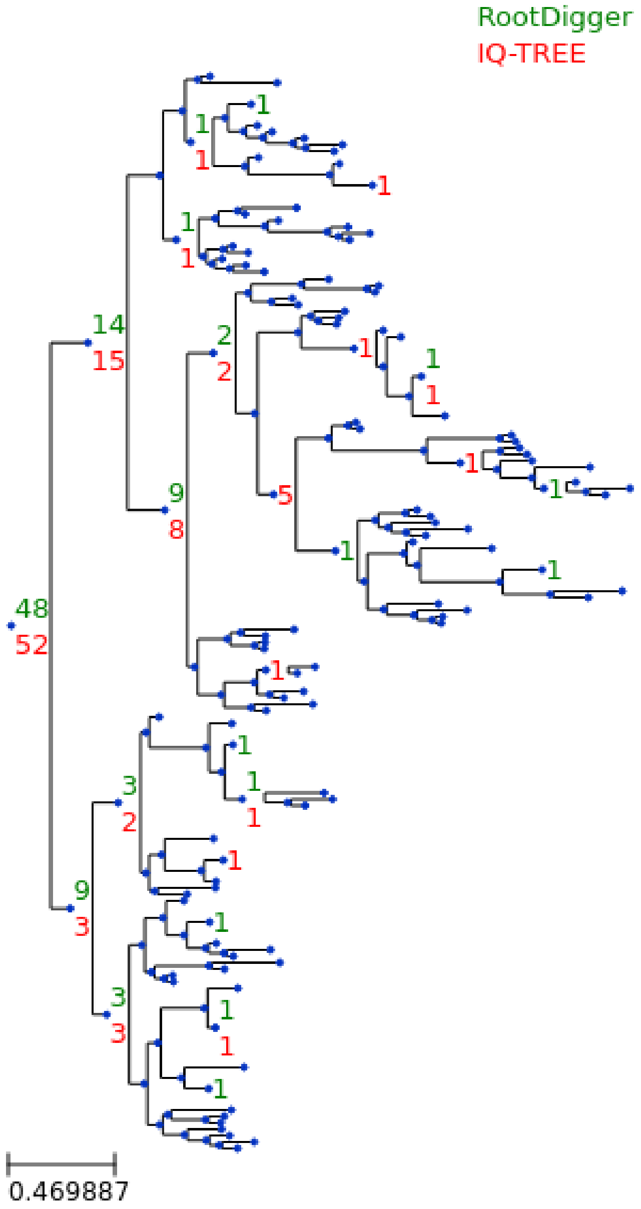


Figure 2: Simulation results for 100 taxa and 1000 sites.

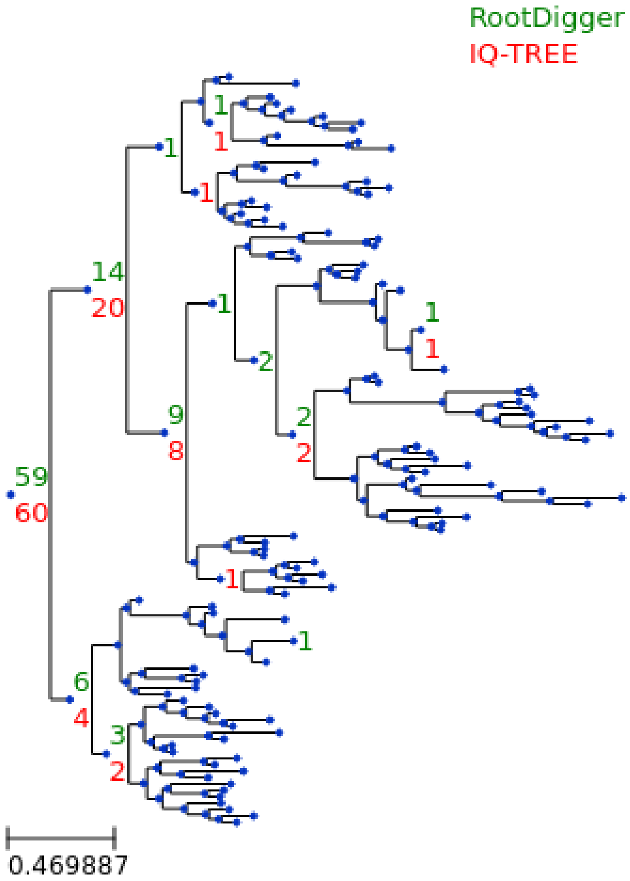


Figure 3: Simulation results for 100 taxa and 8000 sites.

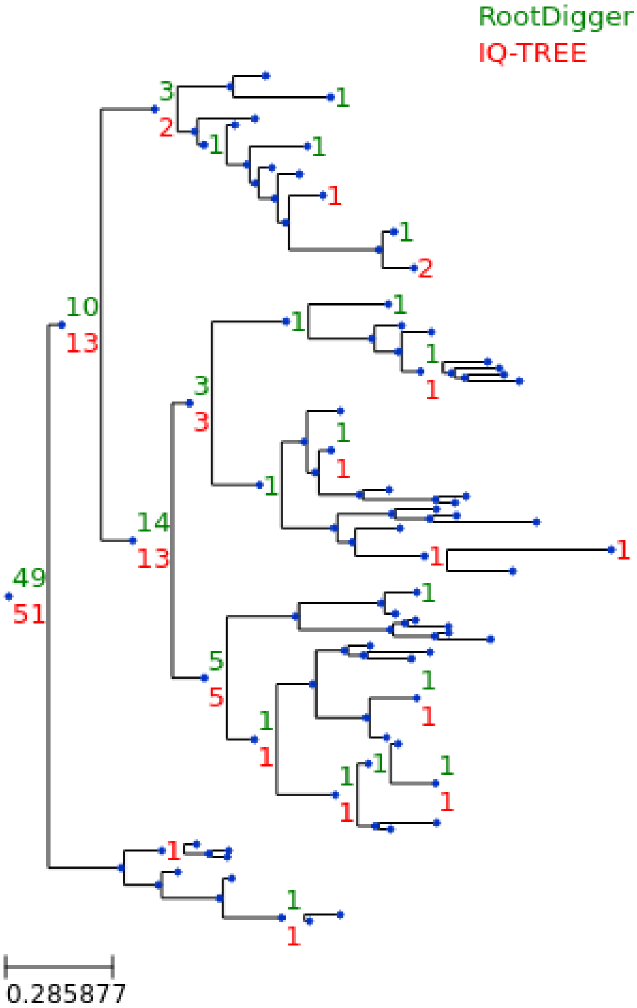


Figure 4: Simulation results for 50 taxa and 1000 sites.

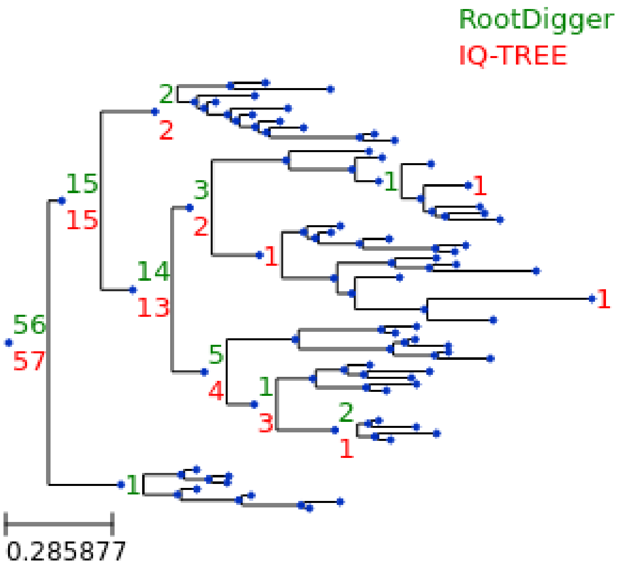


Figure 5: Simulation results for 100 taxa and 8000 sites.

existing, strongly supported outgroup. For each of the empirical datasets, we ran `RootDigger` in exhaustive mode to obtain likelihood weight ratios (LWR) for each branch. Annotations are suppressed for branches with a small LWR (less than 0.0001). The trees with annotated LWR are shown in Figures 6 - 11. We ran the experiments on the datasets with the outgroup included, as well as with the outgroup removed.

There was some preprocessing performed. In order ensure that all sure branch lengths in all datasets were specified in substitutions per site, the branch lengths were re-optimized using RAxML-NG [13] version `0.9.0git`. The original model was used when it was known, otherwise the branch lengths were optimized using GTR+G.

Similar to simulated data, we adjusted the execution time of IQ-TREE by the estimated amount of time spent in branch length optimization. We do this as in simulations, but with the empirical datasets instead.

##### 5.3 Effect of early stopping on result

Finally, we investigated the effect of the early stopping criterion on the final LWR results. To do this, we ran `RootDigger` in exhaustive mode on all empirical datasets with early stopping enabled

Table 1: Table of empirical datasets used for validation.

| Name | Dataset | #Taxa | #Sites | O. Model | Model Used | Source |
|---|---|---|---|---|---|---|
| DS1 | AngiospermsCDS12 | 35 | 864029 | GTR+Γ | GTR + Γ4 | [20] |
| DS2 | AngiospermsCDS | 35 | 1296043 | GTR+Γ | GTR + Γ4 | [20] |
| DS3 | Grasses | 245 | 4973 | GTR+ G + I | GTR + Γ4 | [2] |
| DS4 | Ficus | 200 | 5552 | GTRCAT | GTR + Γ4 | [3] |
| DS5 | SpidersMissingSpecies | 33 | 1097842 | NA[a] | GTR + Γ4 | [14] |
| DS6 | SpidersMitocondrial | 34 | 12479 | NA[a] | GTR + Γ4 | [14] |

[a] The paper states that PartitionFinder was used, but the results were not provided.

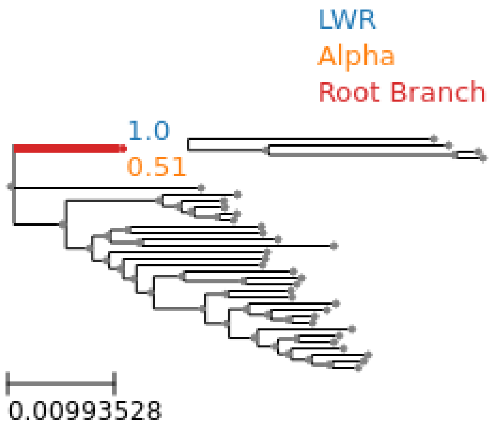


Figure 6: SpidersMissingSpecies dataset analyzed without an outgroup.

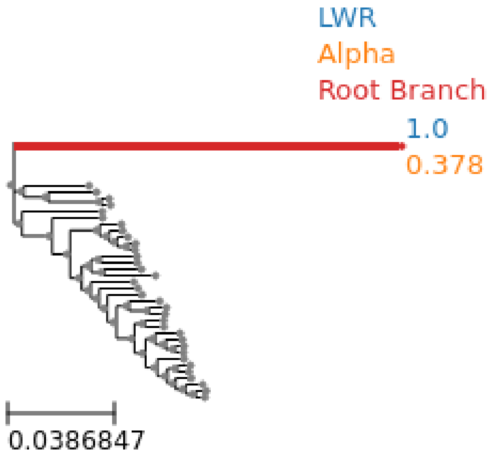


Figure 7: SpidersMissingSpecies dataset analyzed with an outgroup.

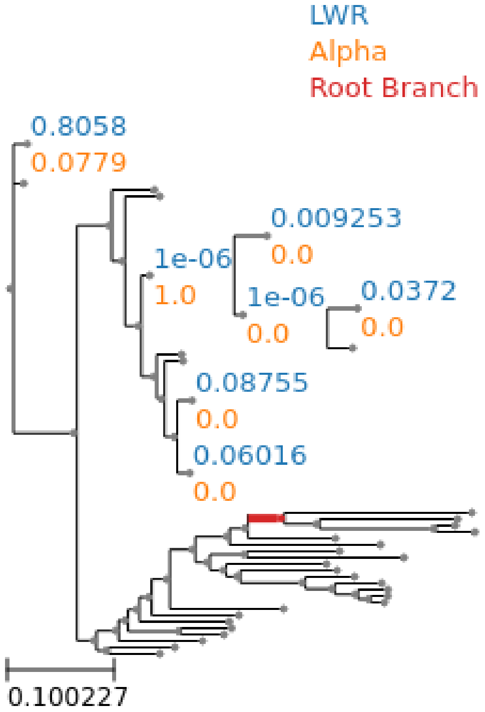


Figure 8: SpidersMitocondrial dataset analyzed without an outgroup.

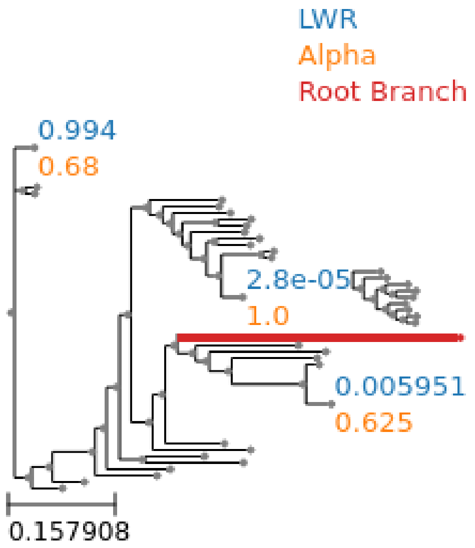


Figure 9: SpidersMitocondrial dataset analyzed with an outgroup.

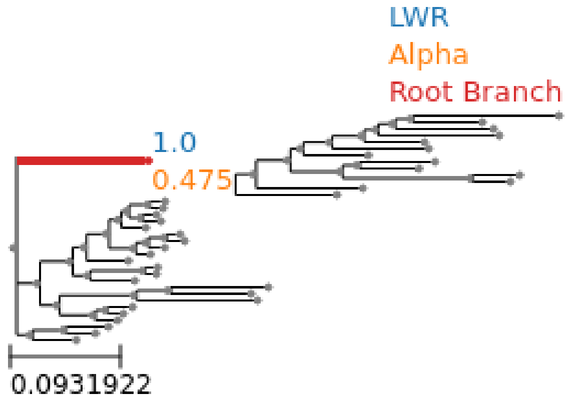


Figure 10: AngiospermsCDS12 dataset analyzed without an outgroup.

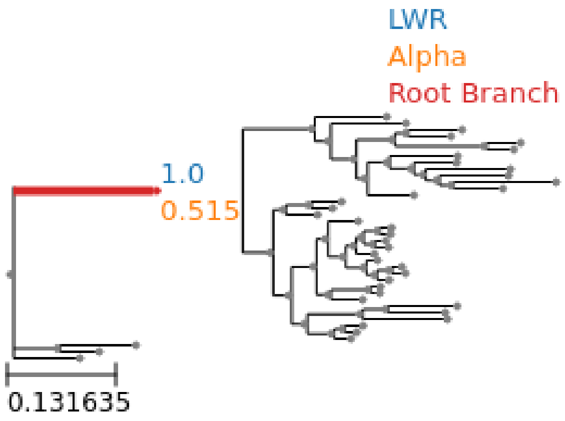


Figure 11: AngiospermsCDS12 dataset analyzed with an outgroup.

and disabled. For most runs, the results with and without early stopping showed no meaningful (difference in LWR less than 0.000001) difference. The dataset that showed the largest difference in LWR is shown in Figure 13. In exchange, the runtime of this dataset with early stopping enabled is about 1.7 times faster.

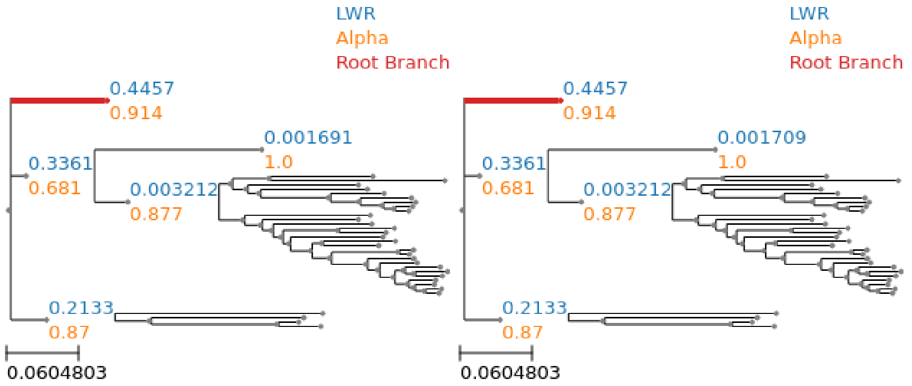


Figure 13: Effect of early stopping on results. Left is with early stopping, Right is without. Dataset is SpidersMitocondrial and has the largest observed difference.

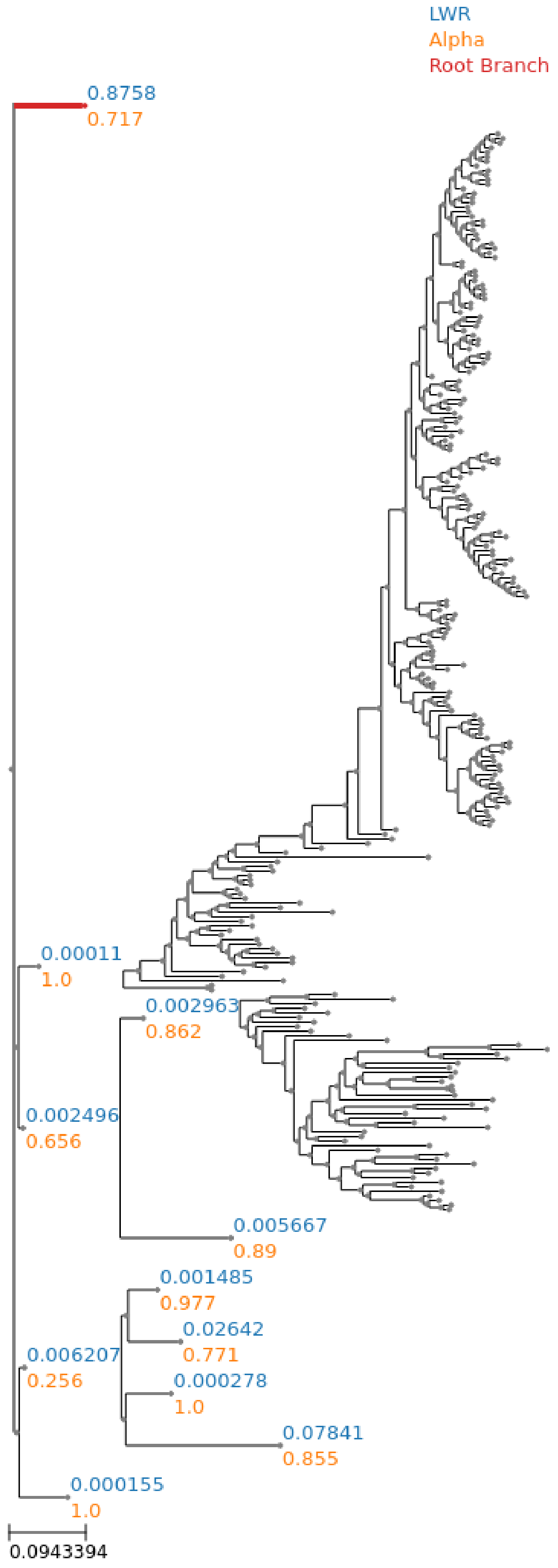




Figure 12: Grasses dataset analyzed without an outgroup.

### 6 Discussion

Compared to IQ-TREE, RootDigger performs competitively, as can be seen in both sides of Figure 1. The results on simulations are mixed, with IQ-TREE performing slightly better in terms of root placement in most scenarios.

RootDigger was faster on all but the smallest datasets. On the smallest datasets, IQ-TREE and RootDigger performed about equal in terms of execution time. Again, it should be noted that IQ-TREE *does* optimize branch lengths, while IQ-TREE does not. RootDigger being faster than IQ-TREE is expected. Nonetheless, IQ-TREE is the only tool that will find the same results as RootDigger with comparable methodology. Therefore, the execution times are only presented here for context.

For empirical datasets RootDigger performed well, with it correctly identifying the root for 3 of the 6 datasets, which is consistent with the results from simulations. The time required to analyze these datasets ranged from approximately 23 minutes (SpidersMitocondrial) to 15 hours (AngiospermsCDS) with early stopping off.

Comparison with IQ-TREE for empirical results is difficult, because IQ-TREE does not produce a distribution of likelihood weight ratios. In order to simulate this to a reasonable degree of confidence, we would need to run IQ-TREE many times in order to produce a comparable distribution of root positions. Given the computational cost of analyzing the largest dataset with IQ-TREE, we chose not to compare with RootDigger on empirical datasets.

Qualitatively, RootDigger offers other advantages: multiple modes of inference (search mode and exhaustive mode), as well as the ability to compute LWR for root positions on a given tree. Again, this can be simulated with many runs of a more standard phylogenetic analysis tool, but the fidelity and speed will be significantly impaired.

RootDigger is substantially faster than IQ-TREE, as is shown when comparing figure 1. Furthermore, the accuracy of root placement of RootDigger is nearly the same as IQ-TREE, as also seen in figure 1. Early stopping proved to be an effective technique to reduce execution times at an extremely marginal cost in accuracy. The largest observed difference on empirical datasets between using early stop mode and is negligent, as it can be seen in figure 13.

The computational improvement of early stop mode on simulated datasets is less apparent than on empirical datasets. Early stop mode on simulated datasets performed at negligibly better (the largest improvement is approximately 2%) compared without early stop mode.

In Huelsenbeck [11], it was shown that the prior probability of a root placement on a sample tree did not have a strong signal when using a non-reversible model of character substitution. While performing our verification of `RootDigger` using empirical data, we found that this was often not the case. For example on the AngiospermsCDS12 dataset (see figure 10), we found a clear signal for the root placement, both with and without the outgroup. Even in cases when the signal was not as strong, for example SpidersMitocondrial (see figure 8), there is a much stronger signal for root placement than Huelsenbeck would suggest we would get with this kind of analysis (which is to say, analysis using a non-reversible model). The one exception to this is the ficus dataset, which showed at least marginal support for the root on nearly all branches of the tree. We suspect that this is due to Huelsenbeck performing the analysis on a 4 taxon tree with the distantly related taxa frog, bird, mouse, and human. By only using 4 distantly related taxa, the rate matrix is less constrained by the data present, which may lead to over fitting the rate matrix. We believe that the methods presented here will typically produce a clear signal for the rooting of a tree.
